# Developmentally timed sensory integration enables efficient larval dispersal

**DOI:** 10.1101/2025.10.07.680954

**Authors:** Marte Lønnum, Madeline M. Schuldt, Jose Davila-Velderrain, Lena van Giesen

## Abstract

Animals integrate internal and external sensory information to adjust movement gait according to the physical constraints imposed by body-environment interactions. Efficient locomotion is especially important in small organisms where the motile phase is ontogenetically restricted. Motile larvae of otherwise sessile cnidarians must disperse and identify a suitable habitat in a restricted timeframe. Here we show that dispersal in *Nematostella vectensis* larvae is accomplished by a constant ciliary sensory-motor system that produces stimulus-induced movement. In contrast, neuro-muscular and sensory systems gradually increase in complexity during development, enabling movement-associated gait control through reafferent matching of external and internal information. Together with ciliary propulsion, the developmentally timed appearance of sensory and neuronal structures endows the animal with the ability to integrate information to shape swimming behavior and achieve dispersal in a timely manner, appropriate to the physical challenges of its specific ecological niche.

## Introduction

Anthozoan cnidarians such as anemones and corals are sessile or semi-sessile marine invertebrates. Their complex lifecycle includes a motile larval stage which serves dispersal, an important feature that ensures the resilience of the species by enhancing genetic diversity and facilitating inhabitation of novel ecological niches^1,2^. During the larval phase, the animals display state-specific behaviors, with planktonic larvae exhibiting dispersal behavior early in development. Animals can disperse and explore the environment passively (e.g. by ocean currents) or actively (through swimming). A second type of behavior occurs when the larvae become competent, a physiological state in which behavior centers around the choice of a permanent settlement site. In this phase, the animals terminate their pelagic residence and start exploring the benthic environment to identify a suitable, and often permanent, habitat. Many aspects of larval dispersal and settlement remain enigmatic. It is not well understood which sensory modalities cnidarian larvae employ to guide swimming or settlement, how such information is processed, how it modifies behavior, and which cellular subsystems are involved.

Anthozoan larvae are diverse in developmental time and anatomy^3^ and consequently larval behavior too is multifaceted and species-dependent, suggesting that both these traits contribute to the adaptation to a specialized ecological niche with its physical peculiarities. Swimming in the larvae can encompass a variety of modes and planes, and some cnidarian larvae even exhibit other complex behaviors such as active predation^4–6^. In addition, within the same species, larval behavior is highly plastic throughout development, potentially mirroring changing goals from dispersal to settlement. Animals have been reported to transition from near motionless to active swimming and later to crawling and gliding or probing the substrate immediately prior to settlement^7–9^.

For both dispersal and settlement, it can be assumed that sensory information is relevant in guiding the larvae. Indeed, both behaviors have been linked to sensory perception in cnidarian planula larvae^6,10–13^. Cnidarian larvae have to date not been found to perform active taxis but rather phobic responses which nevertheless result in specific distribution of the adult polyps with respect to sensory stimuli^13^. During these phobic behaviors, the animals respond to sensory stimuli by modification of their activity with changes in ciliary beating and body shape that result in slow positional changes^6,8,11^. Using phobic responses rather than active tactile swimming is a sensible and energy efficient strategy, considering that, with the size of the larvae and the vastness of the ocean, active swimming towards or away from a stimulus would often exceed the physiological capacity of the animals.

How cnidarian larvae with their small size and relatively simple body plan regulate these diverse behaviors is unclear. The motile planula larvae is polarized and swims forward with its aboral end propelled by cilia, which cover the whole animal, forming metachronal waves that improve the efficiency of locomotion^14,15^. In addition, some anthozoan cnidarians possess a sensory organ, the apical organ with long protruding cilia at the aboral end^16–18^. Furthermore, many other specialized cell types, such as secretory, neurosecretory, putative sensory, and support cells are enriched in the apical region of the larvae, suggesting that this area combines several sensory effector centers that may be involved in guiding specific behaviors during the larval life^18–21^. This idea is also supported by the underlying prominent aboral nerve plexus, of which some parts might be larval specific^16,19,22^. In addition to the above mentioned cell types, muscles become more prevalent and increasingly organized over the course of development^23^, and late-stage planula larvae can contract using their muscular-hydraulic system^24^, a behavior that might be directly modified by sensory input such as light in *A. millepora* larvae^6^.

These recent advances in knowledge about cnidarian behavior are critical to our understanding of the basic biology of this vast group of marine invertebrates that form the foundation of many rich ecosystems. Cnidarians occupy a particularly interesting evolutionary position since they are one of the most basal extant animal lineages with a nervous system that might influence their behavioral repertoire already during early life stages such as the motile planula. While even single-celled eukaryotes can perform sophisticated cilia-mediated behaviors^25,26^ and sponge larvae use both sensory and ciliated cells to modify their swimming pattern in response to light^27^, it is an exciting endeavor to study the exact function of the nervous system and other larval-specific structures, such as the apical organ, to shed light on the complex interplay between cilia and developing nervous and muscular systems.

Here we describe the basic swimming behavior of the *Nematostella vectensis* planula larvae over the course of development from hatching to metamorphosis into a primary polyp. To study if swimming behavior indeed changes over development and reflects life stage specific behavioral interests that are mediated by specialized anatomical features, we analyzed these aspects concomitantly in carefully selected stages. *Nematostella vectensis* is an ideal model system for such a study due to its easy handling and regularly inducible spawns in which fertilization can be precisely timed for stereotypical larval development. *Nematostella* is a burrowing sea anemone, commonly found in shallow estuaries and tidal pools. Its lifecycle includes a motile larval stage with an apical tuft that transitions into a primary polyp (metamorphosis) within a few days (**Fig. 1a**). The sea anemone is a well-developed model organism with a vast variety of tools, such as a sequenced genome, transgenic lines^28^, and single-cell transcriptomics data available for diverse developmental stages^29^.

**Figure 1:**
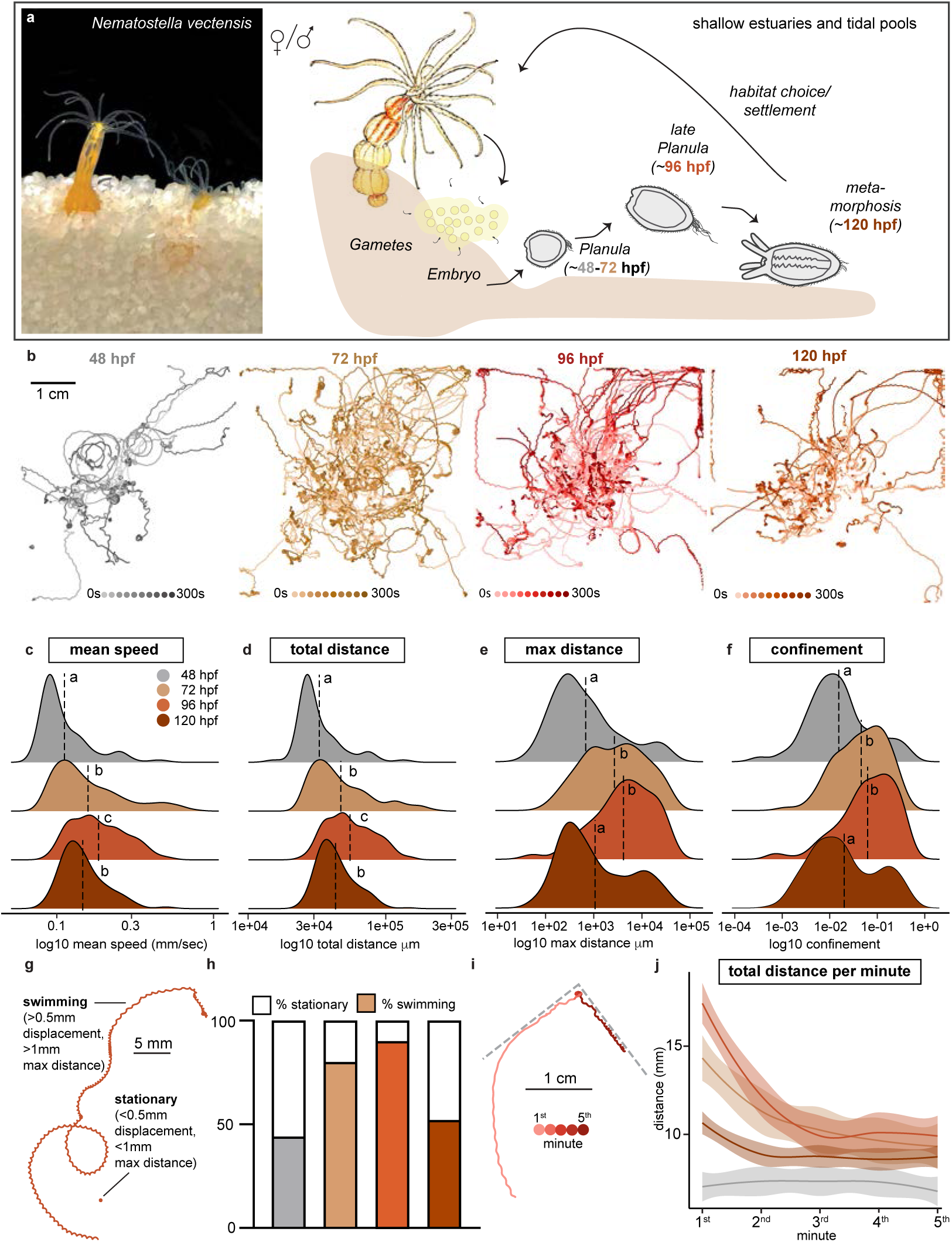
Swimming behavior is changing over the course of larval development. **a)** *Nematostella vectensis* adult (left) and its lifecycle (right). **b)** Overlayed tracks produced during a 5-minute video from four ages of *N. vectensis* larvae showing swimming behavior. 48hpf (n=14, N=123), 72hpf (n=14, N=167), 96hpf (n=12, N=158) and 120hpf (n=16, N=187). **c-f)** Distributions of track mean speed, total distance, maximum distance and confinement ratio. Dotted line at the mean; letters denote statistical significance by Kruskal-Wallis test, x-axes on log10 scale. **g)** The defined thresholds for categorizing larvae as either stationary or moving. **h)** Ratio of moving and stationary larvae per age. **i)** Example trace and **j)** calculations of age-specific average distances the larvae swam per minute. Lines show a smoothed conditional mean line ± SEM, see also supplemental information.

Our data show that swimming behavior changes drastically over the course of development. *Nematostella* planula larvae become faster, and swim both further, and less spatially confined until metamorphosis. This advance in swimming abilities occurs simultaneously with an increase in sensory capabilities, in form of a higher response to mechanical perturbation, a longer apical tuft, and the enhanced expression of sensory receptors, which were found predominantly in the ciliated epithelial cells. Furthermore, while the length of the motile cilia and the body remain relatively stable during the active phase, the animal’s body shape changes both over development and as the animals behave, correlating with specific swimming modes. We find that the nervous system controls the shape-shifting abilities of the late larval stages but has little impact on sensory perception or the ciliary beating per se, suggesting that *Nematostella* larvae use two independent systems to control swimming behavior: a ciliary sensory-motor system and a neuronally controlled muscular contraction system, aimed at optimizing body shape to swimming mode.

## Results

### Swimming behavior is changing over the course of larval development

After preliminary experiments, we chose four ages for behavioral observations at which animals showed clearly distinct behavioral patterns. These timepoints roughly correspond to two phases of the early planula (48 and 72 hours post fertilization (hpf)), the late planula stage (96hpf) and the tentacle bud or metamorphosed stage (120hpf), at our developmental conditions (21°C) (**Fig. 1a**). To assess detailed aspects of behavior at these ages, we developed an assay that allowed us to obtain comprehensive behavioral tracks and the associated metrics under standardized conditions by filming as the animals moved freely for five minutes (**Fig. 1b** and **Supplementary Fig. 1a,b**, see Methods section for details). As previously reported^8^, larvae quickly develop into active swimmers between 48 and 72hpf and become more explorative at 96hpf before mostly ceasing activity around 120hpf (**Fig. 1b**). Our tracked data allowed us to extract detailed aspects of the swimming behavior as the animals developed. For example, we observed that not all metrics changed simultaneously or stereotypically between and within the different ages, indicating that behavioral control is more complex than previously appreciated and may depend on different aspects of the larval development, specifically anatomical characteristics and information processing capacity.

As they age, more larvae swim faster and further, with the average farthest distance (59.5 mm) and highest mean speed (0.2 mm/sec) occurring at 96hpf; however, the individual with the highest total distance (224.5 mm) and mean speed (0.75 mm/sec) was a 72hpf aged individual. The slowest age was 48hpf (mean speed 0.12 mm/sec), which also swam the shortest average distance (35.8 mm). At 120hpf these values drop (mean speed 0.15 mm/sec and total distance 45.5 mm) and reach similar numbers to those observed in 72hpf larvae (**Fig. 1c-d, Supplementary Fig. 1c,e**). The total distance traveled correlates with speed (Range R^2^=0.94-0.98), as these metrics depend on each other (**Fig. 1d, Supplementary Fig. 1j**, see also methods). However, total distance is relative (larvae could have a high total distance value by swimming spatially confined in circles), which would not contribute to an actual euclidean distance, a value that is more relevant for biological phenomena such as dispersal behavior. To determine if larvae also travel further in space and do so more effectively, we measured maximum distance (euclidean distance **Fig. 1e**) and confinement as a measure of efficiency to reach a distant point in space (**Fig. 1f, Supplementary Fig. 1f,i** and methods for calculations). Animals at 48hpf were most confined on average (ratio 0.048), and displayed the lowest maximum distance (3.1 mm). In contrast, 96hpf larvae showed lower confinement (ratio 0.12) and the highest average maximum distance (8.7 mm). Interestingly, when looking at the distribution of these two metrics, we found that 48 and 120hpf old larvae showed distributions with mostly low, but subpopulations of relatively high values. This pattern was not found at 72 and 96hpf, which display a unimodal distribution of similarly high values, indicating that all larvae at these ages and only a small proportion of the youngest and oldest larvae are capable of dispersing efficiently. Intriguingly, these observations parallel the appearance of the apical tuft, a larval specific sensory structure^30^.

As reported by Hand and Uhlinger^8,11^, we also observed that *Nematostella* larvae can switch between stationary and swimming modes (**Fig. 1g**), indicating that active swimming behavior is an energetically costly behavior that the larvae will only display when conditions are advantageous. When quantifying the percentage of stationary versus moving individuals across age, we found that the ratio is shifting in favor of movement in 72 (80%) and 96hpf (89%), while for the youngest and oldest larvae the ratio is approximately 50% (**Fig. 1h, Supplementary Table 1h**).

The range between the minimal and maximal speed values that larvae can achieve is not drastically changing between different age groups (**Supplementary Fig. 1g,h**), suggesting that the physiological requirements for both inactive and fast swimming modes are given throughout the development of the animals. Consequently, the ability or the drive to swim must be actively regulated to explain our observations. Therefore, to understand why the animals display different modes of activity with distinct ages, we looked more closely at the distribution of the activity (total distance) over the time of our 5-minute experiment and discovered that the largest shifts were observed during the first 60 seconds of our recordings (**Fig. 1i,j**). This time point corresponds to the arousal phase that is caused by the mechanical handling of the animals or the transfer to a novel environment, a common feature that has been reported before both in *Nematostella*^8^ and other organisms^31^. This peak in arousal of larvae at 96hpf shows that the animals are more sensitive to external stimuli at this age and coincides with the highest expression of putative sensory receptor proteins, such as TRP channels, some of which have been implicated in mechanosensitive behavior in *Nematostella vectensis*^18,32^ (**Supplementary Fig. 1k,l**).

### Larvae can modify speed and behavioral mode independently

Observing relatively complex locomotor patterns in the older planula larvae, we decided to study aspects of the animal’s swimming trajectories in more detail at 96hpf. We quickly realized that, despite the peak in activity during the first minute on a population level (**Fig. 1i,j**), the swimming trajectories of individual animals are far from stereotypic across the recorded time interval (**Fig. 2a** and **Supplementary Fig. 2a**). Indeed, the example trace in **Fig. 2b** shows in detail that the larvae can control both speed and swimming mode independent of each other, suggesting that these aspects of locomotion are under active control and the larvae are capable of modifying both aspects dynamically. For better resolution and connectivity between the recorded metrics and to exclude behaviors that are caused by interactions with the walls of the larger arena, we tracked swimming behavior under a 2x microscope objective for 15s (for details see Methods section). To understand if certain behaviors are executed only at specific speeds, we divided the mean speed distribution of the larvae into four quartiles (**Fig. 2c**).

**Figure 2:**
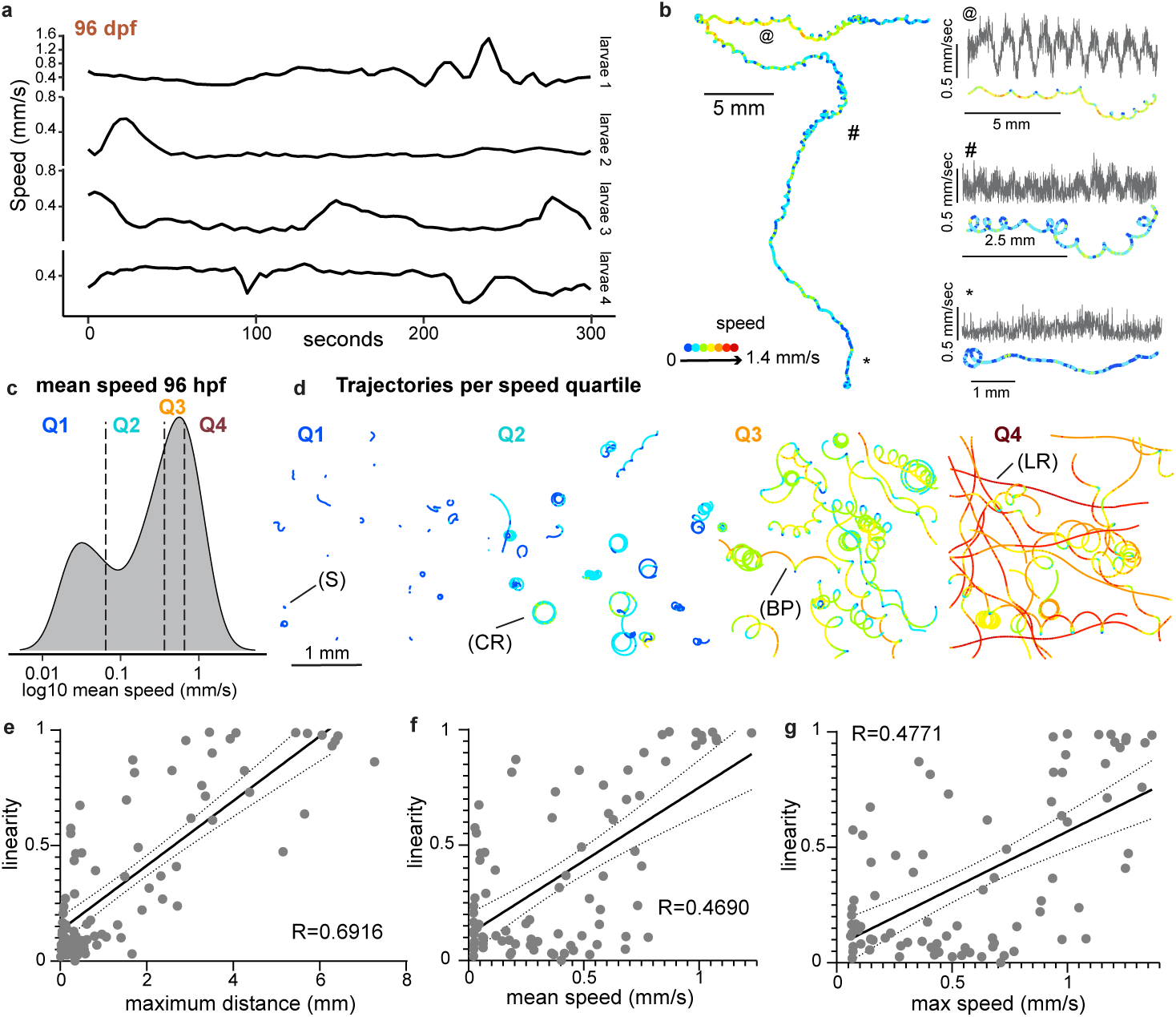
Larvae can modify speed and behavioral mode independently. **a)** A selection of speed profiles from four different larvae. **b)** Example track from a 96hpf larva. Colors show momentary swimming speed. Inserts of three regions as denoted by symbols @, #, * and their corresponding speed profiles. **c)** Distribution of track mean speeds (n=23, N=83) from larvae filmed under a 2x objective, dotted lines denoting the 0.25, 0.50 and 0.75 quartiles (Q1-Q4). **d)** Plots showing larval tracks sorted into the defined quartiles (Q1-Q4). Distinct behavioral modes labeled: stationary (S), linear run (LR), circular reorientation (CR) and bottom probing (BP), colors show momentary swimming speed. Spearman R Correlation of **e)** track mean speed with linearity of forward movement, **f)** track max distance with linearity of forward movement and **g)** track max speed with linearity of forward movement (n=23, N=56).

When plotting the tracks according to their percentile, we observed several swimming modes. Larvae can be stationary (S), perform circular reorientation (CR) or display bottom probing (BP) and linear runs (LR). These modes appeared with different proportions in different quartiles (**Fig. 2d**). There are two major aspects recognized: firstly, larvae achieve higher distances the straighter they swim (**Fig. 2e**, correlation max distance and linearity R^2^=0.6916). Secondly, when looking at the relationships between metrics, we observed that particularly the slowest (stationary) and the fastest behaviors (linear runs) show a strong link with both mean and maximum speed. The stationary mode, which is forming the lower end of the correlation with low mean and maximum speed, is contrasting the linear runs that show the highest mean and maximum speed as well as the farthest distances achieved (**Fig. 2e-g**; correlation linearity and mean speed: R^2^=0.4690, maximum distance: R^2^=0.6916 and maximum speed R^2^=0.4771). These data show that stationary behavior and linear runs are highly correlated with the animal’s speed mode while other behaviors can be executed at a variety of speeds. Together with the data from the larger arena, it becomes clear that while animals can perform distinct behavioral sequences, overall more larvae of an older age are swimming linearly and therefore faster (both displaying higher maximum and mean speeds) and thereby disperse more efficiently to a distant point in space, suggesting that throughout development, the animals gain better control over their movements and that the drive to disperse upon sensory stimulation is particularly high at 96hpf.

### Body shape changes over development affect locomotion behavior

Given our observations, we next asked which aspects of the larvae change over the course of development that would enable the animals to swim more linearly and therefore disperse faster and more efficiently when agitated. We first measured the body shape of the motile stages of the planula larvae (48-96hpf) (**Fig. 3a**). Anatomical elongation can be quantified using the body axis ratio, a measurement where higher ratios between the length and the width indicate a more elongated shape. Body axis ratio, as measured from whole-body images, was found to increase with age (mean ratio 48hpf: 1.172, 72hpf: 1.333, 96hpf: 1.379) (**Fig. 3b, Supplementary Fig. 3a,b, Supplementary Table 3b**). Since it has been reported that cnidarian larvae routinely perform body shape shifts during active behavior^6,9,33^, we wanted to investigate whether freely swimming animals of the same age show variations in body shape, and how these variations correlate with locomotor activity. The microscopy objective tracks obtained from **Fig. 2** enabled us to extract swimming metrics and a simultaneous measurement of the body shape (**Fig. 3c**). Since the body shape can change due to both contractile activity as well as reorientation in z (**Fig. 3d**, see bottom probing trace wine red as example), we focused our correlative analysis on animals that remained in a stable z position over the course of the 15 seconds. Indeed, longer animals swim, on average, faster, as indicated by a positive correlation between axis ratio and mean speed that was found in 96hpf larvae (**Fig. 3e**, R^2^=0.5117). On the other hand, the total size of the larvae (measured as area) is neither correlated with speed (R^2^=-0.0611) nor with elongation (R^2^=0.05427, **Supplementary Fig. 3c-d**), showing that body elongation but not total body size is contributing to increased swimming speeds.

**Figure 3:**
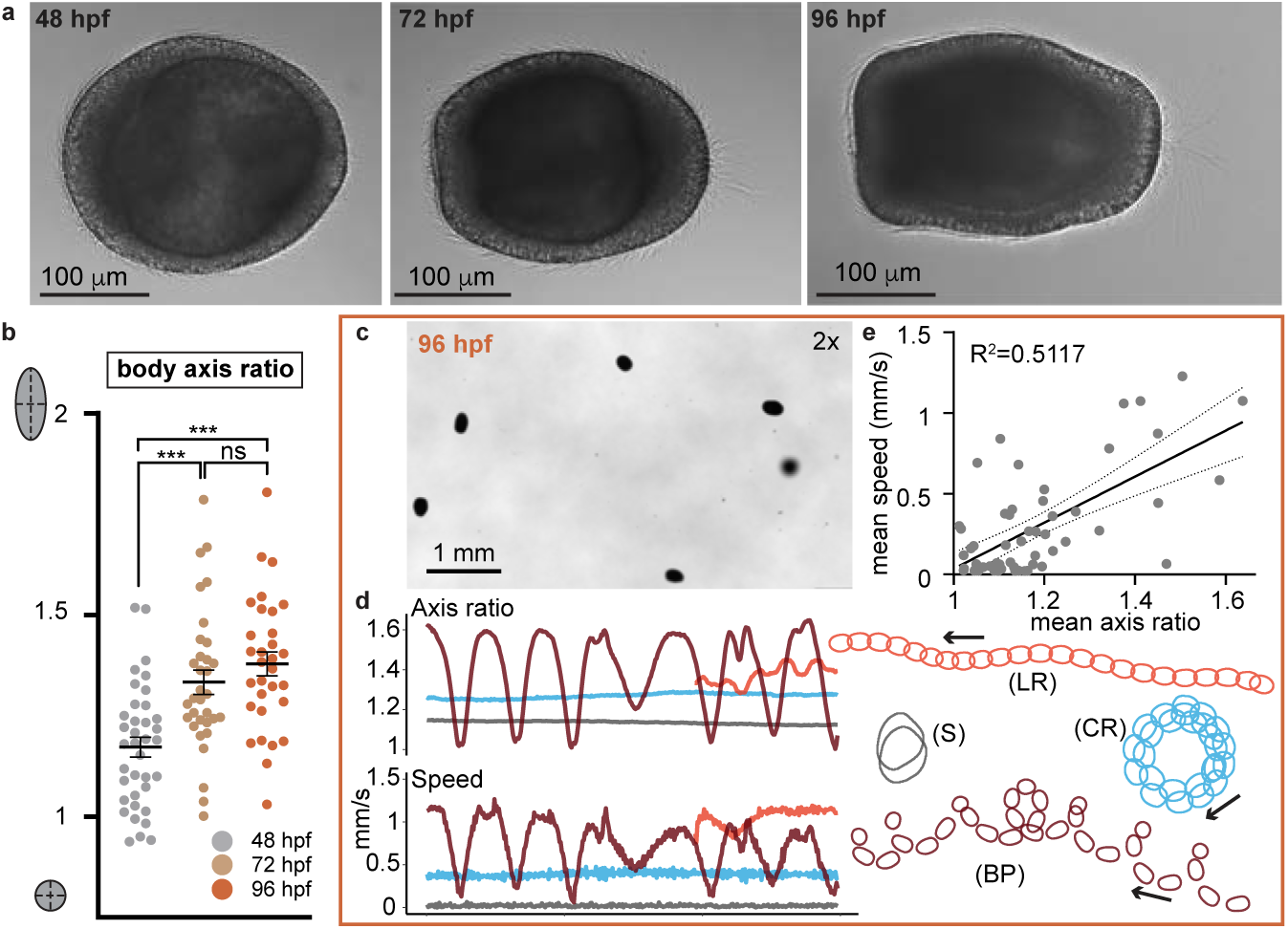
Body shape changes during development and locomotion behavior. **a)** Representative images of motile larvae over the course development at 48, 72 and 96hpf at 20x. **b)** Body axis ratio of larvae at rest significantly increases after 48hpf. Lines at mean ± SEM; statistical significance by Tukey’s multiple comparisons. **c)** View of freely swimming larvae through the 2x objective. **d)** Axis ratio and speed profiles of four different larvae performing four distinct behavioral modes; stationary (S), linear run (LR), circular reorientation (CR) and bottom probing (BP). **e)** Spearman R Correlation between mean body axis ratio and mean swimming speed (n=23, N=56).

How exactly shape changes of the larvae influence its swimming behavior and efficiency is unclear. One possibility is that the ciliary tuft of the apical organ is the reason for the elongation of the aboral area. We observed that larvae in linear runs and bottom probing behavior often show some constriction towards the apical tuft and that the tuft is bundled and pointing straight ahead (**Supplementary Fig. 3e**). This could enhance the polarity of the larvae either for better steering if the tuft serves a paddle function as suggested in^34^ or for further sensory reach.

### Cilia change in length but not basal beating frequency

Along this line of thought, we next investigated which aspects of larval ciliation are changing over the course of development. In cnidarian planula larvae, cilia act both as sensory and motor units and can most likely modify swimming pattern and behavioral complexity through both functions^16,35,36^. We therefore investigated the overall length of the epithelial ciliation in the aboral, side, and oral region, (**Fig. 4a,e**) as well as their beating frequency (**Fig. 4b-d**) and the length of the sensory organ cilia, the tuft (**Fig 4f**), to understand more about how ciliary details might contribute to the enhanced sensitivity and better steering abilities in older larvae.

**Figure 4:**
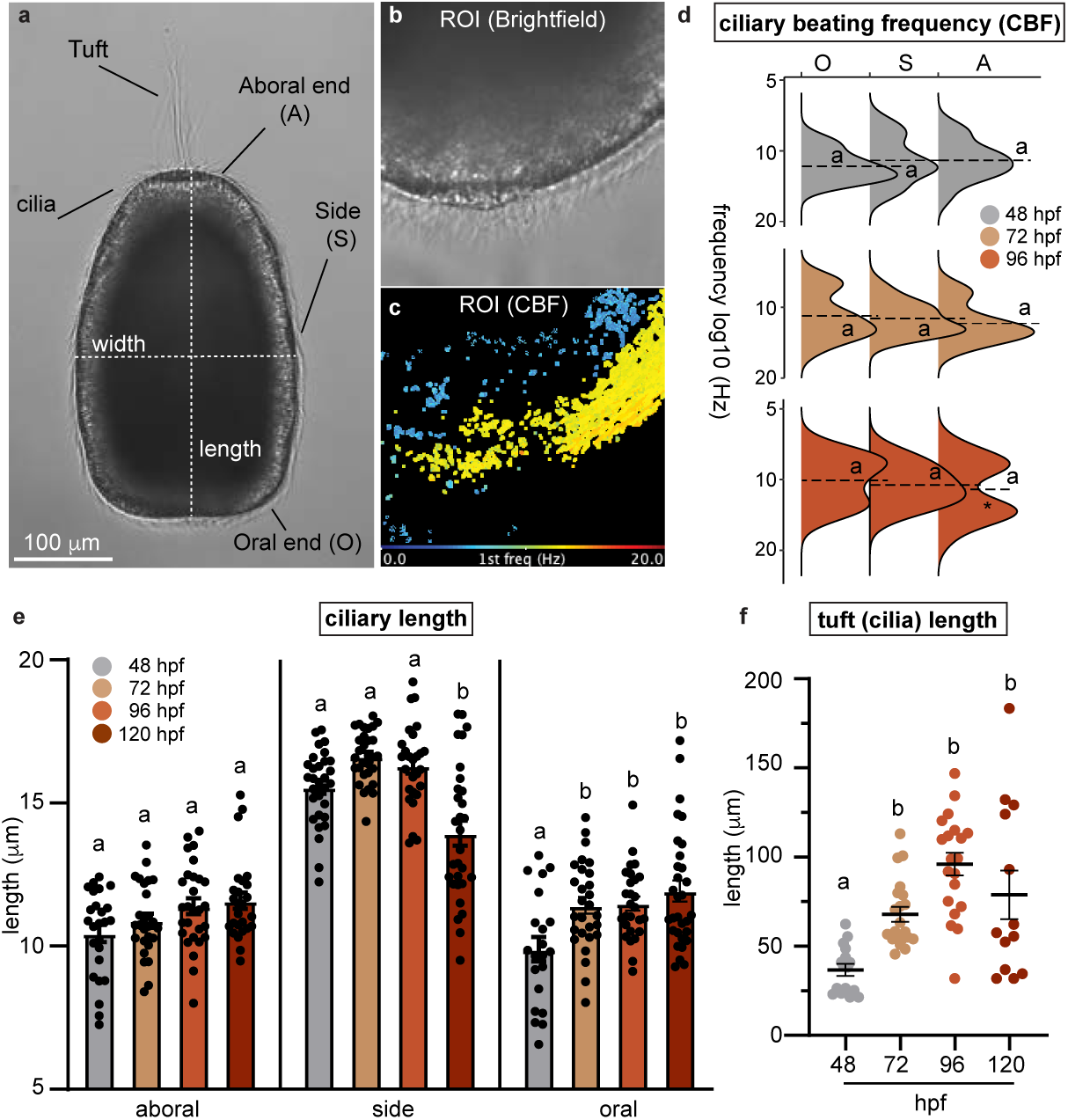
Cilia beating frequency and length during development. **a)** Representative image of a 96hpf larva at 20x with regions of measurement denoted. **b)** Image of region-of-interest (ROI) taken of a high-speed microscopy video taken for CBF analysis. **c)** Frequency map depicting peak frequency for each pixel. **d)** Cilia beat at frequencies between 7 and 18 Hz with no significant differences. Dashed line represents the means; asterisk (*) denotes a significant deviation from unimodality. **e)** Cilia at the apical region of the body do not significantly change in length with age, whereas cilia along the side of the body decrease in length after 96hpf and cilia at the oral end increase in length after 48hpf. **f)** Tuft (cilia) length shows a trend of increasing length after 48hpf. Lines at mean ± SEM. Letters denote statistically significant differences in means by Šídák’s multiple comparisons (ciliary length) and Dunn’s multiple comparisons (tuft cilia length).

Our measurements in restrained larvae show that the ciliary beating frequency (CBF) ranges between 7 and 18 Hz for all ages and body regions (**Fig. 4d** and **Supplementary Fig. 4a**), with no significant differences between the means. This observation shows that there is no overall increase in beating frequency as the animals age, and that larvae at all ages might be able to actively modify the CBF between set boundaries. Larvae at 96hpf show a bimodal distribution of beating frequencies in the aboral region. It is possible that this sharp distribution points to an enhanced control over the beating at this age. Our setup does however not permit statements about CBF changes during swimming behavior, which might be more relevant to determine its effect on the speed in freely moving animals.

Other aspects of ciliation, as for example increased ciliary length, might correspond to a larger power stroke when beating, increasing the force generated to propel the larval body through the water. We measured the length of cilia across the larval body over development (48-120hpf), and when comparing average values for the three regions within the same age we found that cilia in the side region are significantly longer (range of means 13.94-16.63 µm) than those in the apical (10.45-11.58 µm) and oral (9.89-11.93 µm) region for all ages (**Supplementary table 4e**). When comparing ciliary length within the same region across different ages, we find three primary features (**Fig. 4e**). Firstly, there are no significant differences in apical ciliary length across ages. Secondly, 120hpf larvae have significantly shorter side cilia compared to 48-96hpf animals (mean 48hpf: 15.55 µm, 72hpf: 16.62 µm, 96hpf: 16.31 µm, 120hpf: 13.94 µm), hinting towards a possible role in locomotion of this subgroup of cilia. Finally, 72-120hpf larvae have significantly longer oral cilia compared to freshly hatched larvae (mean 48hpf: 9.89 µm, 72hpf: 11.42 µm, 96hpf: 11.49 µm, 120hpf: 11.93 µm), possibly in preparation for a more prominent role of the oral end in feeding in the metamorphosed animals (**Fig. 4e**).

In addition, we measured the length of the cilia comprising the tuft of the apical organ and found significantly longer tuft cilia in animals older than 48hpf (**Fig. 4f**, mean 48hpf: 36.81 µm, 72hpf: 67.91 µm, 96hpf: 96.07 µm, 120hpf: 78.86 µm). Longer cilia increase both the physical reach and can house a greater number of receptors due to a larger membrane surface, leading to enhanced sensitivity for sensory stimuli, providing better spatial information that could facilitate more agile or faster swimming behavior^37,38^. This is especially important considering the putative sensory role of the specialized cilia of the apical tuft. As the tuft cilia lengthen, not only does the larva experience an improved sensory capability but also an enhanced polarity. A sensory role of motile cilia along the side of the body would further increase environmental perception to the whole body of the animal, leading to better flow sensing and improved proprioception, thereby facilitating complex swimming behaviors^39,40^. At 120hpf, apical tuft cilia show a wide spread of overall lengths and high variability (SEM of 13.55 µm, more than double that of any other age) (**Fig. 4f, Supplementary Table 4f**), indicating the onset of the loss of the apical organ as previously reported^20^ and indicative of a decreasing need for this sensory structure when the larvae become non-motile. The timing of the loss of this structure coincides with the loss of motility but occurs before settlement. This reinforces the notion that the apical tuft is required for processing sensory information related to dispersal and swimming behavior, rather than settlement, and possibly forms a structure that aids in steering^17,34^.

### Cilia and neurons play distinct roles in swimming behavior

Since the function of the larval ciliation, particularly the apical tuft, the nervous system, and their interplay in locomotion remain poorly understood in *Nematostella* larvae, we were keen on describing these structures concomitantly. By using the same timepoints as the behavioral experiments, we hoped to deduct how anatomical differences correlate with the observed behavioral changes. To this end, we performed immunohistochemistry of larvae aged 48hpf-120hpf with anti-DsRed in the transgenic *Elav::mOrange* line that selectively labels neurons (**Fig. 5a**)^41^ and with anti-acetylated tubulin to show the ciliary structures (**Fig 5b**). In addition to the histology, we re-analyzed single-cell transcriptomic data at relevant developmental stages^29^ to understand how cellular and molecular components endow the animal with distinct functional subsystems that enable a more controlled swimming pattern over time (**Fig. 5c-e** and **Supplementary Fig. 5a**).

**Figure 5:**
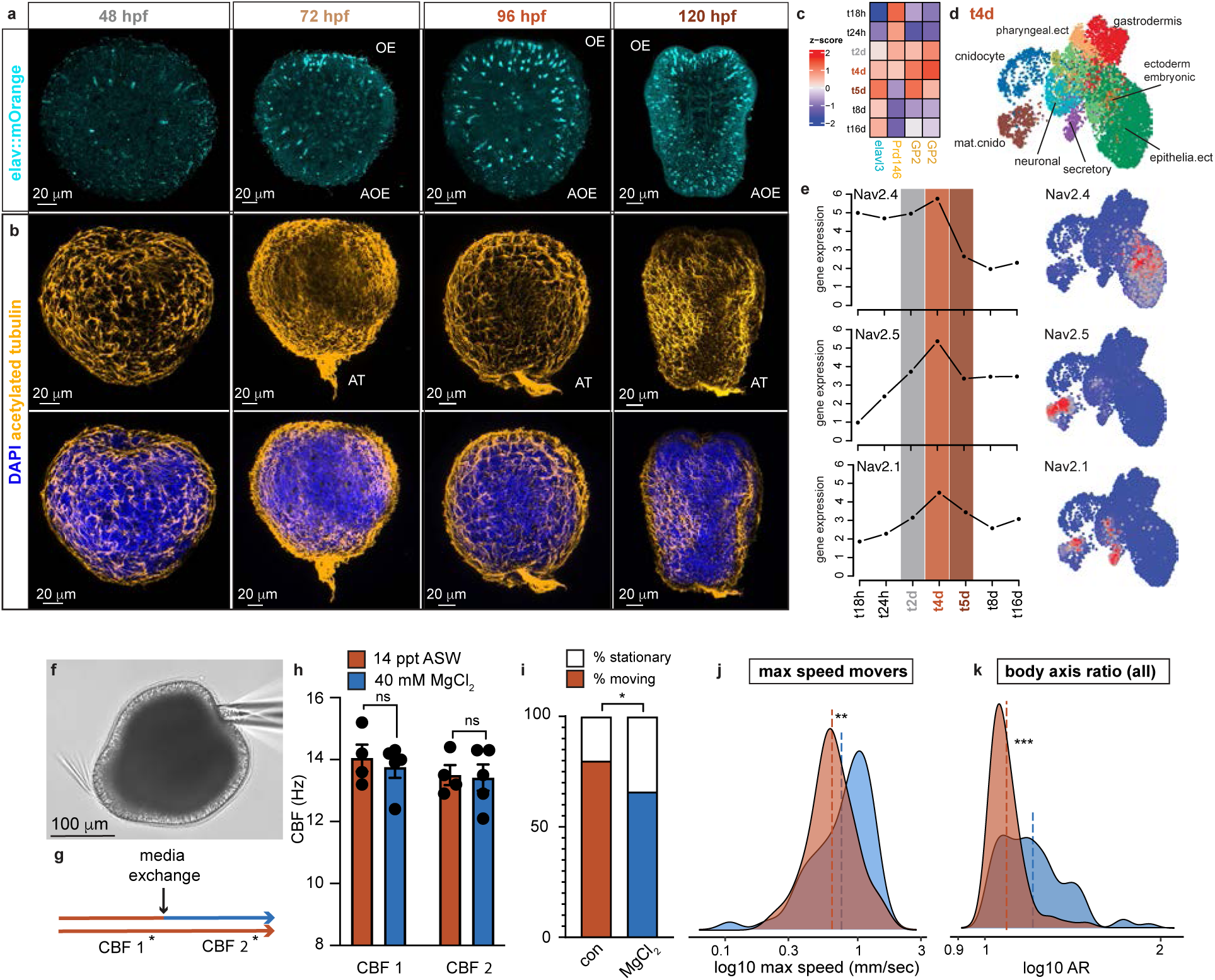
Nervous system and cilia are distinct subsystems involved in swimming behavior. **a)** *Elav::mOrange* larvae stained with dsRed between 48-120hpf. **b)** Ciliation of larvae between 48-120hpf visualized using acetylated tubulin (yellow) and DAPI for nuclear stain (dark blue), (OE= oral end, AOE= aboral end, AT= apical tuft) **c)** Expression profiles of marker genes. Data show relative average expression values across developmental time. **d)** Two-dimensional representation of single-cell transcriptional landscape at age td4 (∼96hpf). Data points represent single cells labeled by cell groups reported in ^29^. **e)** Developmental average expression profiles (left) and 2D projection of single-cell expression values (right) of genes encoding Nav ion channels. **f)** Representative image of a 96hpf pipette-attached larva at 20x. **g)** Schematic of pipette-assay protocol. **h)** Comparison of CBFs of larvae in the control vs treatment conditions at both CBF 1 and CBF 2 video timepoints revealed no significant differences in CBF when exposed to MgCl_2_ for 5 minutes. Lines at mean ± SEM (control n=4, MgCl_2_ n=5). **i)** Ratios of moving and stationary larvae for the control and MgCl_2_ treated larvae. Two tailed two-proportion Z-test *p= 0.039. **j)** Distributions of maximum speeds between the control and MgCl_2_ treated larvae (moving animals only). Line at the mean, **p=0.005 by Mann-Whitney test. **k)** Distributions of axis ratios between the control and MgCl_2_ treated larvae. Line at the mean, ***p<0.0001 by Mann-Whitney test.

We found that the larvae show an increasing number of neurons, as previously reported^42,43^ with a small number of ectodermal sensory neurons in the early planula stage and larger numbers in both ecto- and endoderm in the late planula. Finally, a clearly visible nerve net is found by the time animals are developing prominent tentacle buds (**Fig. 5a**). Remarkably, we observed that the largest number of sensory neurons in the aboral area appears only in the 120hpf stage (**Fig. 5a**, right), suggesting that older larvae have an increased need for sampling substrates. *Nematostella* larvae indeed metamorphose before they settle and can reattach for a long period of the early polyp stage^8^. In agreement with the literature and our measurements (**Fig. 4f**), the 48hpf larvae rarely possess an apical tuft. This structure appears later and remains prominent up until 5dpf, when it becomes less dense and some larvae were observed to have already lost the tuft. Interestingly, when looking in the single-cell data set from^29^ using markers for neurons (elavl3), the apical organ (Prd146) and the ectodermal epithelium (GP2) from^20^, we found that, while the neuronal marker is increasing in agreement with our immunohistological observations, both the apical organ ciliated cells (Prd146) and some ectodermal cells labeled with GP2 seem to be largely constrained to the motile phase of the larvae (**Fig. 5c,d** and **Supplementary Fig. 5a**). These observations are consistent with the expression of Nav2.4 as a proxy for the functional capacity of the ciliated epithelial cells. Nav2.4 is strongly expressed in younger larval ages and disappears almost completely after the animals seize swimming activity (**Fig. 5e**). This sodium channel is exclusively found in the larval ectoderm and apical organ cells^44^ and (**Fig. 5e** right panel), suggestive of a role in excitability in these ectodermal cells (e.g. ciliary beating, contractility). Contrasting to Nav2.4, strong expression of Nav2.1 and 2.5, sodium channels that are characteristic of other excitable cells, such as neurons and cnidocytes, is observed only in later larval stages. These data suggest that the larvae loose molecular and potentially also cellular components associated with ciliary locomotion. All together, these observations make it likely that individual systems play a role at different stages in the animal’s life and, in addition, serve distinct functions with respect to ontogenetically restricted behaviors.

To further test this hypothesis, we blocked synaptic transmission using magnesium chloride (MgCl_2_)^45^ and observed how the inhibition of the nervous system affects ciliary beating and swimming behavior in 96hpf larvae, the motile stage with the most developed nervous system. Baseline ciliary beating as measured in larvae fixed by a pipette (**Fig. 5f**) did not change when MgCl_2_ was applied as compared with control animals, which remained in normal media (**Fig. 5g,h**). Cilia in the larvae, therefore, beat without neuronal input and are, at least during basal conditions, not influenced by neuronal activity. Ciliated cells might create their own sensory-motor units, regulating beating frequency intrinsically in response to external information, such as mechanical or light stimuli, rather than being influenced by neuronal input. This finding is supported by previous observations of sensory receptors being expressed directly in epithelial cells in *Nematostella* planula larvae^18,35^ and by our expression analysis, which finds a large proportion of the highly expressed TRP channels at 96hpf in epithelial cells (**Supplementary Fig. 5b**).

When magnesium chloride was applied to freely moving animals, we observed that, despite the lack of an obvious change in CBF, fewer animals were among the actively swimming proportion (66% moving) compared to the control (80% moving) (**Fig. 5i**, for the definition of moving vs stationary see materials and methods). While the reduction of motile animals was significant, the majority of the planula larvae remained activated through the arousal by handling. This result strengthens the theory of cilia as independent sensory-motor units and shows that a fully functioning nervous system is not basally required for swimming but rather constitutes an additional layer of locomotor control.

To further investigate what this role could be, we looked more carefully at the swimming and body shape differences between control and magnesium-treated animals. Magnesium-treated animals showed substantial changes in both parameters (**Fig. 5g,h**). Actively swimming larvae (movers only) (**Fig 5g**) swam faster (median mean speed 0.88 mm/sec) as compared to the control (median mean speed 0.63 mm/sec). When plotting the distribution of the speed quartiles of all animals (**Supplementary Fig. 5c,d**), we found that magnesium-treated animals were essentially locked in two speed modes: either stationary (Q1) or in high speed (Q4) and only a few larvae were found in the intermediate speed modes. These results indicate that neurons are contributing to the animal’s capacity for gait switching. Movement gait is an acquired feature, and this concept is consistent with our findings. *Nematostella* larvae do not show changes in the range between maximum and minimum speed but rather appear to learn to utilize their body more efficiently over developmental time. Locomotion gait is mediated by matching internal and external sensory information to reach optimal swimming patterns. Therefore, larval swimming gait would include proprioceptive responses and appropriate adjustment of the body shape to a given external sensory environment. Indeed, when plotting the axis ratio of the animals, it became evident that the inhibition of synaptic transmission leads to a disconnect between body shape and speed (**Supplementary Fig. 5e**), as well as extreme shape deformations such as elongation and crumbled body shapes (**Supplementary Fig. 5g**). Swimming gait is important for maximal locomotor efficiency and endows the animal with the capacity to fine-tune its behavior in agreement with external and internally collected sensory information. Reorganizing the body shape through muscular contractions can influence ciliary orientation and consequently lead to speed modifications induced by flow field changes. These changes will in turn influence proprioceptive signaling, helping the animal to assess its behavior and orient itself with respect to its environment, facilitating fine-tuned behavioral modes^40^.

## Discussion

In the present study, we report, for the first time, detailed aspects of swimming behavior of the *Nematostella vectensis* planula larvae across systematically chosen developmental timepoints and examine its relationship to concomitantly appearing cellular and molecular features. Larval behavior is changing over time, leading to optimized dispersal through the integration of external and internal sensory information that enables the animal to “learn” to utilize their form in a more efficient way. While some aspects remain relatively stable over development, such as minimum and maximum speed, length of motile cilia, and their beating frequency, the larvae instead shift on the scale of possibilities towards more linear trajectories and faster speeds through the polarization and sharpening of sensory functions. The advance of the sensory systems allows animals to be more active and “sensitive” in response to external stimuli and, additionally, establishes a reafference system based on neuronally mediated body shape contractions. This sensory integration makes *Nematostella* larvae versatile swimmers with a rich repertoire of behavioral modes.

For an animal with the size of the *Nematostella* larvae, swimming in aqueous medium is energetically extremely costly. Indeed, ciliated epithelial cells are metabolically highly active^36^ and the ciliary length might be perfectly adapted to swimming efficiency in small organisms^46^. Given that a helical swimming pattern is stereotyped across several marine species, some aspects of swimming are potentially also confined by physical laws that determine the maximal efficiency of movement and optimize flow fields^14,26,47,48^. While *Nematostella* swimming speed (0-3mm/sec in still water) and its body size may not endow the animal with the capacity to resist larger currents, their swimming ability is well adapted to facilitate dispersal and habitat selection within their natural ecological niche of tidal pools and marshes, where water may remain still or experience incremental increases. Responding to sensory stimuli by increasing both activity and fine-tuning behavior, reflected in the ability to control swimming modes, facilitates dispersal and enables the larvae to invest their efforts with maximal efficiency. This could occur for example either during favorable conditions, if a current is sensed that might allow the animal to disperse further away from the colony with less effort, or if conditions become unfavorable such as might be experienced by high intensity of light. Sensory information with a negative valence could induce a shift to highly linear swimming, enabling the larva to move quickly away to other areas. Higher activity in larvae has been observed after mechanical stimulation or in response to short wavelengths of light^11,31^. In contrast, responses to stimuli can also induce the opposite response, including drastic body contraction and the cessation of various types of activity^6,13^. How exactly the animal responds to a certain type of sensory information will depend both on the developmental stage and the ecological niche of the animal, making these aspects important considerations in future studies. For example, constraining dispersal time both during development and also acutely, as observed for *Nematostella* planula larvae, is an adaptive behavior for an anemone that lives in relatively restricted area^49^, where large dispersal distances could lead to a loss of surviving offspring to the vast ocean currents.

Furthermore, sensory perception will also include self-sensing in the form of proprioception and flow sensing, leading to optimization in swimming gait^40^. The ability to actively and effectively utilize the set of swimming modes available to the animals, and to transition between those modes at appropriate timepoints, seems to be governed by an intrinsic mechanism that “turns on” at a certain developmental stage. This increase in sensory abilities co-occurs with increasing numbers of TRP channels in epithelial cells and concomitant elongation of the apical tuft. Being able to access the external and internal sensory information allows the animal to be activated through appropriate stimulation and to optimize swimming gait to reach their goals in the ontogenetically available time. Larvae that have gained “access” to such swimming modes could expand this efficiency to other aspects of their behavior; whereby sensory information might induce a shift to bottom-probing with which the larva can gather more information about the sediment.

How cnidarian larvae sense their environment is not well understood. Recent investigations into the cellular and molecular architecture of different cnidarian planula have however opened the possibility for detailed investigations of such aspects^19,20,29,50^. In agreement with previous observations that have shown that the apical organ is not required for the larvae-polyp transition^20,21^ and is lost in some animals prior to settlement^17^, our data suggest a sensory role of the apical tuft mainly in swimming behavior. Larvae display larger activation through mechanical input and more linear trajectories at ages that show a prominent tuft, while, at competency, when most animals begin the search for a suitable habitat, the tuft is degrading or lost. Several studies have found sensory receptors expressed in the apical organ^18,35^ and, together with reports documenting the apical tuft to move as a unit during swimming with a sweeping movement^16,17^, it is likely to possess a tactile-sensory or stirring purpose, providing clear polarization of the elongated planula larvae that creates a sensory directionality to guide swimming movement. The increase in tuft length with age might further increase sensory perception by allowing more receptors to be housed in an individual cilium, which would increase the sensitivity of this organ^51^. Which other sensory modalities *Nematostella* larvae use and how the ciliated cells communicate among each other and with the nervous system will be of great interest in future investigations.

In addition to the tuft cilia, our analysis reveals that larvae might not just lose the tuft, but possibly a larger fraction of sensory-motile cilia that enable swimming behavior in the planula larvae. These epithelial cells show expression of functional ion channels only during the motile stage of the larvae, such as Nav2.4 and TRP channels, strongly suggesting that these cells might serve a sensory-motor purpose too. Indeed, previous reports have found sensory-motor cells in cnidarian planula larvae^36^ and it is well known that most cilia can sense external stimuli. How and if these ciliated cells and their cognate molecular machinery are involved in the larvae’s swimming is unclear at this time.

A second aspect of improved swimming control is the development of the nervous system. When and how the first nervous systems appeared, and what their role was, is under much active debate. It has been suggested that the nervous system in ciliated metazoan larvae first appeared for signal amplification and direct sensory motor connections^52^. Our data show that this is a reasonable suggestion, but also that the situation might be more complex. Firstly, we find that many sensory receptors and other “neuronal” ion channels such as voltage gated sodium channels are expressed in the ciliated epithelium, possibly in some form of larval specific sensory-motor unit. This ciliated sensory-motor unit appears to be in large parts independent of the nervous system, since blocking synaptic transmission only partially impairs swimming and mostly affects the fine-tuning of behaviors that are mediated by muscular contractions. In other cnidarian larvae it has been observed that the nervous system is absent or rudimentary and that ciliary sensory motor units might be the only means of steering the animal^36^, a feature that is maintained from more basal ciliated larvae^52^.

We suggest here that the function of the nervous system is the optimized orientation of the sensory organ and possibly other ciliary structures, such as the swimming cilia, which may be controlled by muscular contractions and relaxations. Such active reorientation of the ciliated epithelium might be used to position ciliary rows along the proper body axes, producing more efficient thrust irrespective of ciliary beating frequency. Previous studies have shown that body shape and ciliation can be modified for dispersal efficiency^53^ and that the generation of coherently directed flow requires cilia to be optimally oriented^54^. The body shape and the resulting sensory experience additionally influence the swimming ability and connect appropriate speeds with specialized behaviors. Shape-shifting has been reported in several cnidarian larvae^6,9,33^ and further analysis might shed more light on how exactly the complex body shape patterns observed here serve specialized functions in the larva.

Understanding the details of behavior in cnidarian planula larvae is of paramount interest for several reasons. Basic biological principles about how basal metazoans integrate information from sensory-motor units that are more ancient (cilia) and neuro-muscular systems unique to animals, can advance our understanding of the roles these two subsystems play and how their interaction enables more complex behaviors suited to the habitat and lifestyle of the animal. Furthermore, understanding aspects of this critical phase in the larva’s life is important to solve questions related to the current climate crisis and loss of biodiversity. To protect ecosystems, we must understand them. *Nematostella vectensis* has served as a pioneering model into the biology of cnidarians and it is urgently required that we develop more tools to understand the fundamental aspects of cnidarian biology. In the present study, we systematically characterized the swimming behavior of the planula larvae as the animals develop to describe some key aspects of their behavior. Future studies should venture into cellular and molecular aspects of sensory perception as well as other types of behaviors. Comparative studies can additionally further our knowledge on how the planula larvae disperse and subsequently find a permanent settlement site with respect to their ecosystem and its physical characteristics.

## Methods

### Animals

*Nematostella vectensis* wild type and *Elav1::mOrange* were a gift from Fabian Rentzsch (University of Bergen) and maintained after^55^. In brief, sea anemones were kept in glass Pyrex dishes filled with 14 ppt artificial sea water (ASW, Coral Pro Salt, Red Sea Aquarium System, R11220) and fed 2-3 days a week with freshly hatched brine shrimp nauplii (*Artemia sp.*). Animals that were on a spawning cycle were kept at 18°C in darkness. The anemones were induced to spawn regularly and after fertilization and development at RT (21°C), larvae and polyps were used at indicated ages for specific experiments.

### Behavior

#### Horizontal swimming

Swimming behavior was performed with *Nematostella* larvae aged 48, 72, 96- and polyps at 120 hours post fertilization (hpf). In each experiment 12 larvae of a specific age were transferred into the center of a custom-made chamber (10 x 50 x 50mm) containing 5 mL of 14 ppt ASW at RT and filmed immediately after transfer at 24 fps for 5 minutes using a Nikon Z50 DX 16-50 camera and a Nikkor MC 105/2.8 S lens. To illuminate the chamber evenly, four 8-RGB LED NeoPixel Sticks (Adafruit, product ID 1426) were placed at 0°, 90°, 180° and 270° at a 18mm distance from the outer edge of the chamber (**Supplementary Fig. 1a**) and controlled through an Arduino Mega 2560 Rev3 and Arduino IDE 2.2.1 software (RGB settings: 2, 2, 2, which equals 3,0904E^13^ photons/cm^2^/sec measured from the center of the arena).

#### Swimming and body shape changes in 2x

All animals in this experiment were 96hpf old and experiments were performed at RT. The larvae were placed in a Low-Profile Open Diamond Bath Imaging Chamber (RC-26GLP, Warner Instruments) filled with 750 µL of 14 ppt ASW and placed on a Nikon Eclipse Ti2-U inverted microscope with a 2x objective lens. Two to three 15-second timelapse videos at 25 fps were obtained, using a Hamamatsu Orca camera (model C13440-20CU, light intensity 3.0. MS 2 with diffuser), to record swimming behavior and larval body shape, and the video with the lowest number of individuals meeting the exclusion criterions was retained for analysis.

To describe swimming behavior and body shape changes to neuronal inhibition, larvae were preincubated with either 40mM MgCl_2_ in 14 ppt ASW or 14 ppt ASW alone (control) for 5 minutes in a 6-well plate, transferred into the Diamond Bath Imaging chamber and immediately filmed using the same settings as described above. In this data set, due to the consideration of drug exposure time, the first video was always used for analysis. Each replicate was given a separate well for incubation.

To measure larval body size (Area and Axis Ratio) the “Analyse particles” tool in Fiji ImageJ was used on a binary version of the videos. Tracks and Edge data from TrackMate as well as the body size measurements from “Analyse particles” in FIJI were exported. Tracks from larvae that were touching each other, out of focus or partially out of frame were excluded from further analysis.

#### Behavioral analysis (Particle tracking)

Videos from the horizontal assay and 2x swimming assay were analyzed using Fiji ImageJ^56^, and the plugin TrackMate^57,58^ for particle tracking. The Hessian detector and Simple LAP tracker were employed to create particle tracks. Particle track measurements (metrics) (Supplementary Fig. 3a) included (I) total track distance (the sum of distances between all coordinates, where d_i,i+1_ is the distance from one spot to the next spot in the track), (II) track mean speed (the mean of all momentary velocities within one track), (III) maximum speed (the momentary velocity with the highest value), (IV) minimum speed, (V) track displacement (the distance in a straight line between the first (d_i_) and last coordinate (d_x_) of one track), (VI) maximum distance (the longest distance between any two coordinates in one track), (VIII) linearity of forward progression (a relative measurement where (VII) straight line speed is divided by (II) mean speed) and (VIII) confinement ratio (a relative measurement where displacement is divided by the total distance). Data were analyzed in GraphPad Prism for Windows (GraphPad software) and R Version 4.3.1 (https://www.R-project.org/).

I. Total distance travelled = Sum_i,j_ (d _i,j_)
II. Mean speed = Mean(d _i,i+1_ × s^-1^)
III. Maximum speed = Max(d _i,i+1_ × s^-1^)
IV. Minimum speed = Min(d _i,i+1_ × s^-1^)
V. Displacement = Δd = d_x_ - d_i_
VI. Max distance travelled = Max_i,j_ (d _i,j_)
VII. Straight line speed = Displacement/time
VIII. Linearity of forward progression = Straight line speed/mean speed
IX. Confinement ratio = Displacement/total distance travelled

### Ciliary beating frequency

All experiments in this section were performed at RT in 0.2 μM filtered 14 ppt ASW.

#### Wax assay

To quantify ciliary beating frequency (CBF) larvae were placed on a glass coverslip fitted with a dental wax channel (∼250 μm wide) and allowed to adjust for 2 minutes in ambient light followed by 30 seconds under microscope light before imaging. Videos of free-moving cilia of immobilized larvae were recorded with a 40x objective (Nikon Eclipse Ti2-U inverted microscope with a Hamamatsu Orca-flash 4.0 camera (model C13440-20CU) from three body regions (apical, side, and oral) and acquired at 150 frames per second (fps), with a 3 millisecond (ms) exposure and 18% light intensity, maximally 90 seconds from the end of the adjustment period. Full body images were also taken for body metric quantification.

#### Pipette assay

To quantify CBF over longer time periods and when exposed to 40mM MgCl_2_ in 14 ppt ASW, larvae were tethered via gentle suction using a borosilicate glass micropipette pulled by a P-97 Flaming/Brown micropipette puller (Sutter Instruments). Pipettes were broken off at the tip and fire-polished to give a rounded edge and small diameter. Perfusion of filtered ASW began immediately after tethering. Images and videos were taken for CBF analysis after 5 minutes, immediately after which the perfused liquid was either maintained or switched to the 40mM MgCl_2_ solution and the media in the chamber carefully replaced with the experimental liquid to ensure a consistent environment. A second set of images and videos were taken after 5 minutes. Videos were acquired at 150 fps with a 2 ms exposure and 3.2% light intensity.

#### CBF and body metrics analysis

Analysis of recordings was conducted in FIJI Image J^56^. Videos were aligned to correct for x-y drift using the plugin MultiStackReg^59^ and analyzed for ciliary beating frequency using the plugin FreQ^60^. Cilia length and tuft cilia length were measured from the point at which the cilium emerges from the larval body to its tip. Each point represents the average of one individual larva, comprising an average of 8-12 cilia. Length and width of larvae were measured, and body axis ratio calculated by dividing length by width. Data were analyzed in GraphPad Prism for Windows (GraphPad software, www.graphpad.com) and R Version 4.5.0 (https://www.R-project.org/).

#### Imaging

Larvae of the desired age were selected and placed individually on a glass coverslip in 2.0 μL 0.2 μM filtered 14 ppt ASW in a light squish-prep to restrict larval movement in the z-plane. Larvae were then imaged with a 20x objective (Nikon Eclipse Ti2-U inverted microscope with a Hamamatsu Orca-flash 4.0 camera (model C13440-20CU)).

### Immunohistochemistry

IHC was performed after ^41,61,62^. Briefly, larvae were relaxed in 2.43% MgCl_2_ in 14 ppt ASW for 15 min, fixed in 4% cold paraformaldehyde in PBS for 1h on a rocker with ice. Larvae were then incubated in 10% DMSO in PBS 20 minutes at RT, and with 2% hydrogen peroxide in PBS for 15 min at RT. Larvae were washed with PBST (TritonX 0.3%), 10-15 times, and incubated in 5% Natural Goat Serum (NGS, Merck, G9023) in PBST for 1h at RT. Incubation in primary antibody (ABs) (Mouse anti acetylated Tubulin 1:500 (Merck, T6793) and Rabbit anti-DsRed 1:100 (Takara Bio, 632496)) in 5% NGS followed at 4°C for 60-70h. After subsequent washes with PBST (10-15 times), and 1h incubation with 5% NGS, secondary ABs (all 1:200: Goat anti-Mouse Cy3 (Abcam, AB97035) and Goat anti-Rabbit Alexa fluor 647 (Thermo Fisher scientific, A-21245)) were applied overnight on a rocker at 4°C. Samples were subsequently washed with PBST and DAPI 1:1000 in PBS (Thermo Fisher scientific, D1306) was applied for 60 minutes at RT. Finally, the samples were washed 3-5 times in PBS, mounted in ProLong™ Glass Antifade Mountant (Thermo Fisher scientific, P36980), and imaged with a Zeiss 800 Airyscan Confocal microscope Images were processed using FIJI and Adobe Photoshop.

### Statistical Analysis

Statistical significance *p<0.05, **p<0.01, ***p<0.001

Different letters denote significantly different mean values, where p<0.05.

#### Horizontal swimming

Track data (metrics) and Edge data (speed information) were extracted from TrackMate. Tracks from individuals too close to another individual, where TrackMate lost the larvae and stitching was not possible were excluded from the analysis. Data were checked for normality (D’Agostino & Pearson test, and Anderson-Darling test) and found non-normally distributed for all metrics. Therefore, a Kruskal-Wallis test with Dunn’s multiple comparisons test was used as a non-parametric alternative to ANOVA. For any correlation between metrics, we used the non-parametric Spearman r correlations.

For moving vs stationary larvae definition: The threshold for being characterized as a stationary larva was displacement < 0.5mm, and maximum distance < 1.0mm. The age-specific average distance the larvae swam per minute was calculated and a smoothed conditional mean line using the geom_smooth() function in R was generated.

#### 2x swimming

A Spearman r correlation was performed in Graphpad Prism between mean speed and mean axis ratio, and mean area only for larvae that remained in the same plane during the experiment. Correlations between any metrics were performed by using non-parametric Spearman r correlations. For the mean speed distribution data, the four quartiles (0.25, 0.50, 0.75, 1.0) were calculated in R and using these values the tracks of larvae were plotted according to which quartile they belonged to. Stationary larvae were defined as larvae with max distance <333μm and displacement <310μm. Differences between ratios of stationary to moving for MgCl_2_ data was calculated by Two tailed two-proportion Z-test. Differences in mean speed, max speed, and mean axis ratio between control and MgCl_2_ treated larvae were calculated using a Mann-Whitney test.

#### Wax assay

All data were checked for normality using both the D’Agostino & Pearson test and the Anderson-Darling test. To test for differences in both cilia and body metrics over development, the following analyses were performed: Differences in CBF within age-groups was assessed using a Kruskal-Wallis test with Dunn’s multiple comparisons. For CBF data within region-groups, a mixed-effects analysis with Šídák’s multiple comparisons was used. Hartigan’s Dip Test was applied to assess multimodality. A small number of groups within the ciliary length data were determined to be non-normal, however, due to sample size and differing variabilities, an ANOVA was determined to be the best method for statistical analysis. Therefore, for cilia length data within region-groups, a one-way ANOVA with Šídák’s multiple comparisons was performed; testing for differences in cilia length within age-group was also done using a mixed-effects analysis with Šídák’s multiple comparisons (results not described here). For body axis ratio, a one-way ANOVA with Tukey’s multiple comparisons was performed, and tuft (cilia) length was compared using a Kruskal-Wallis test with Dunn’s multiple comparisons.

#### Pipette assay

CBF data in pipette-attached larvae were determined to be non-parametric. To compare the control and treatment conditions for each CBF timepoint individually, a Mann-Whitney test was done. To compare CBF 1 in the control condition with both CBF 1 and CBF 2 in the MgCl_2_ (treatment) condition, a Kruskal-Wallis test with Dunn’s multiple comparisons was performed (results not described here).

### Single-cell transcriptomic analysis

Whole-body single-cell RNA sequencing data of *Nematostella vectensis* was obtained from ^29^. The developmental subset of the cell atlas corresponding to stages t18h to t16d was extracted for further analysis. Transcriptional pseudobulk profiles were estimated per age group using the normalized sum of total read counts over cells of each group. Read counts were normalized to counts per million (CPM) and log-transformed in base 2. Pseudobulk expression values were used to estimate temporal expression trends of individual gene markers and the average behavior of groups of genes. The cellular landscape at age td4 (equals t96h in **Fig. 5** and **Supplementary Fig. 5**) was analyzed using the computational framework ACTIONet^63^. Briefly, a low-rank approximation of the normalized count matrix is obtained using the single value decomposition (SVD). This reduced data representation is subsequently decomposed using archetypal analysis to define a low-dimensional representation for each individual cell that is useful in measuring cell similarity and building a cell manifold capturing cell relationships in transcriptomic space. The network is projected in 2D coordinates for visualization using the UMAP algorithm. All these steps were implemented using the function *runACTIONet* with default parameter values. To visualize gene expression levels across the cellular landscape, a network diffusion algorithm was used over the cell network to smooth genewise sparse expression values. Network diffusion is implemented in the ACTIONet’s *networkDiffusion* function.

### Identification of putative sensory receptor ion channel genes

To systematically identify putative voltage-gated ion channel proteins, the PFAM family HMM models Ion_trans (PF00520) and Ion_trans_2 (PF07885) were used as query for searching against the reference proteome of *Nematostella vectensis* version NV2 (wein_nvec200_tcsv2) (https://simrbase.stowers.org/starletseaanemone). The resulting protein candidates were then annotated with the best-matching human protein. Best human protein hits were determined by querying each putative channel sequence against the complete set of human protein-coding genes using the HMMER function phmmer. Candidates matching both Ion_trans/Ion_trans_2 families and human transient receptor potential (TRP) channels were considered as putative sensory receptor genes. All sequence searches were performed using the hmmsearch program of the HMMER software package for sequence analysis v3.4 (http://hmmer.org/).

## ACKNOWLEDGEMENTS

We thank Fabain Rentzsch (UiB) for gifting the WT and *Elav1::mOrange* animals and members of the van Giesen Lab for help with *Nematostella* maintenance. We are grateful for feedback on the manuscript from members of the van Giesen Lab and Fabian Rentzsch. This research was supported by the ERC StG “EnvIronchannel” (101076516) granted to L.v.G.

## AUTHOR CONTRIBUTIONS

ML performed behavioral analysis, microscopy, and immunohistochemistry; MMS performed imaging experiments, ciliary beating, and body measurements. JDV performed computational analysis. All authors conceptualized the study, designed and created the figures, performed statistical analysis, and wrote the manuscript.

## Supplementary material

**Figure S1:**
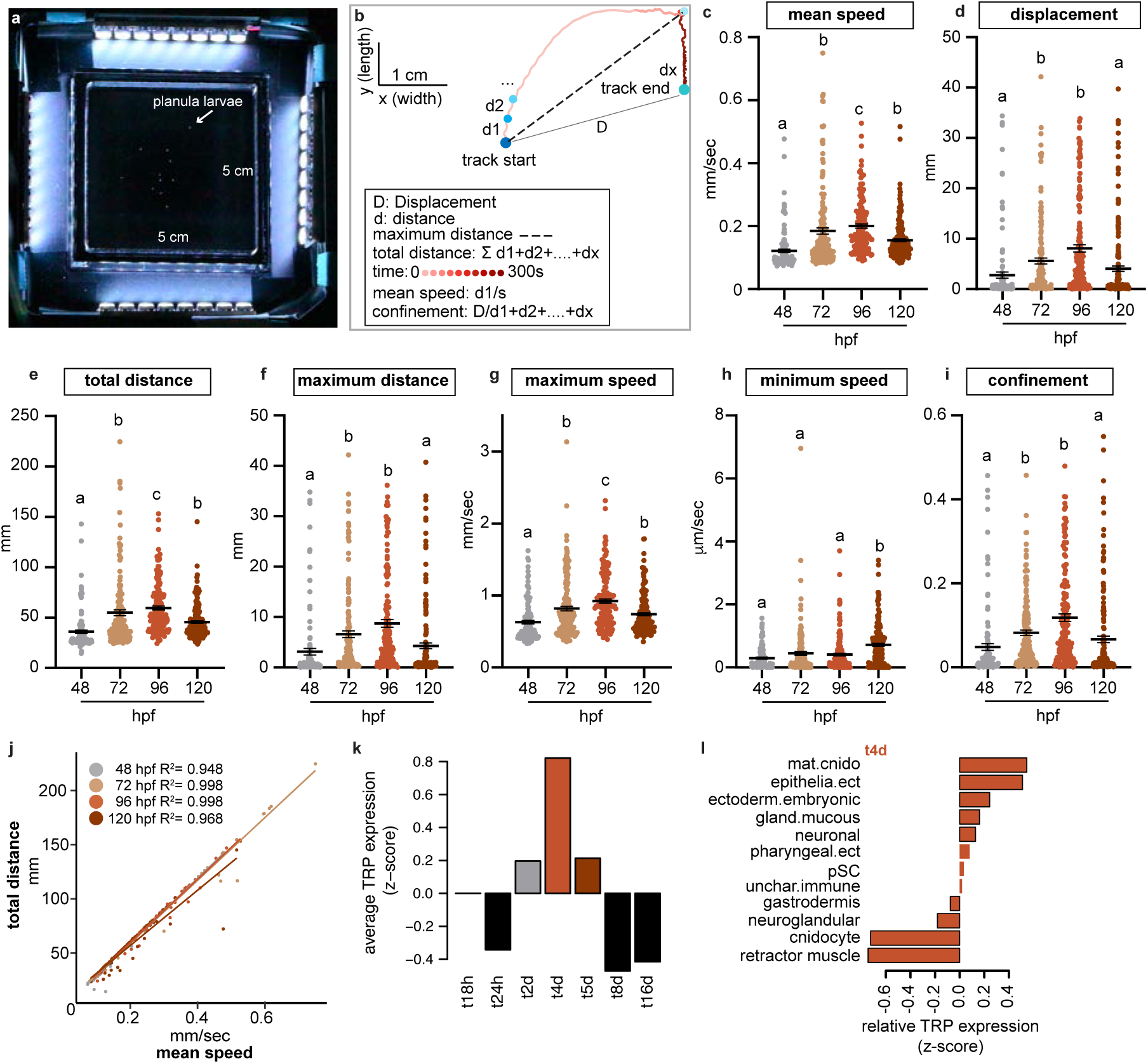
Tracking set up with metrics and expression of TRP channels. **a)** The behavioral arena used for horizontal swimming experiments. **b**) Explanations of various metrics calculated by the tracking software. **c**) Mean speed. **d**) Displacement. **e**) Total distance. **f**) Maximum distance. **g**) Maximum speed. **h**) Minimum speed. **i**) Confinement ratio. **c-i**) Each point represents one individual larva; lines represent the mean value ± SEM; letters denote statistical significance by Kruskal-Wallis test. **j**) Correlation between mean speed and total distance for each age. **k**) Average relative expression profile of putative TRP channels across developmental time. Data shows the average value of relative expression (z-scaled) profiles of putative TRP channel encoding genes (n=50 genes). **l**) Average relative expression profile of putative TRP channels across cell groups at developmental time td4. Data shows similar values as those in k) but estimated by cell group at the time with highest expression (td4).

**Figure S2:**
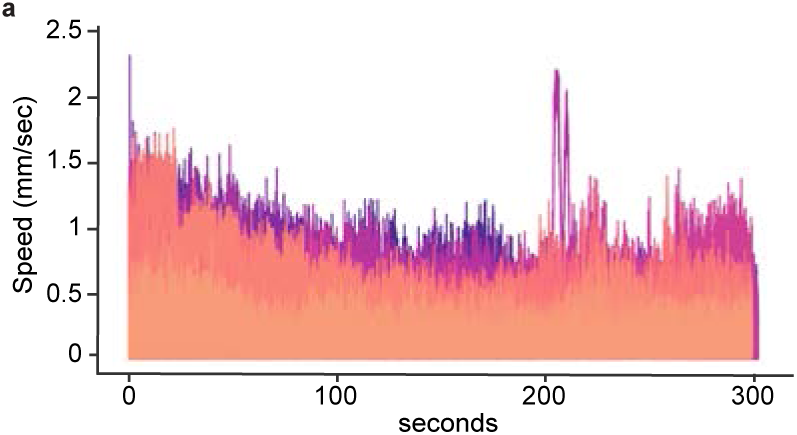
All individual speed tracks for 96hpf larvae. **a)** An overlay of speed profiles (speed per second) of individual larvae at 96hpf. Colors represent different larval IDs.

**Figure S3:**
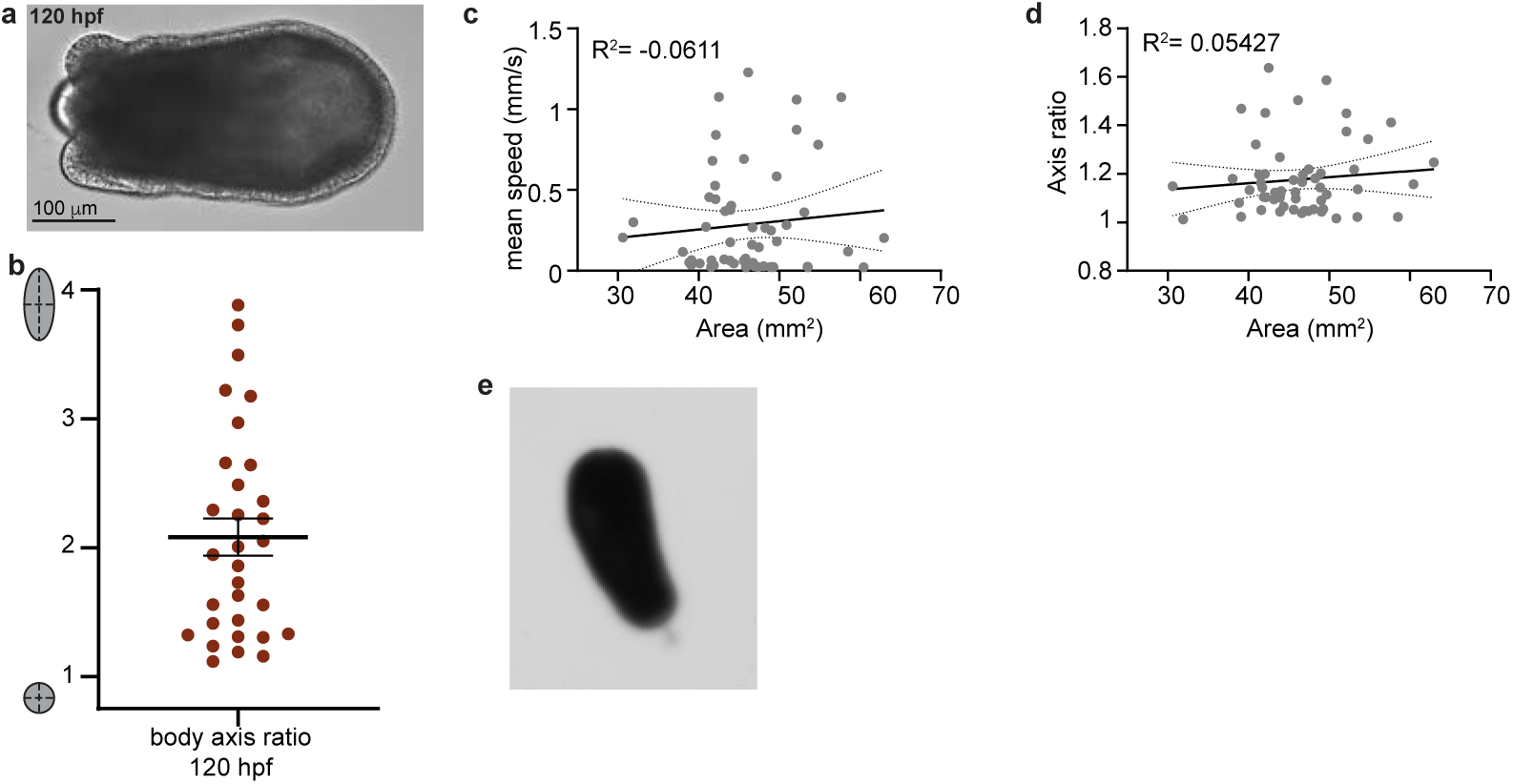
Body shape measurements and correlations. **a)** Representative image of a 120hpf larva in the tentacle bud stage at 20x. **b)** Body axis ratio of larvae at 120hpf. Line at mean ± SEM (n=31). **c)** Correlation between body area (mm2) and mean speed (mm/sec) for 96hpf larvae (n=23, N=56). **d)** correlation between body area (mm2) and axis ratio for 96hpf larvae (n=23, N=56). **e)** Representative still-image of a 96hpf larvae actively swimming with a visible tuft, taken from 2x assay video.

**Figure S4:**
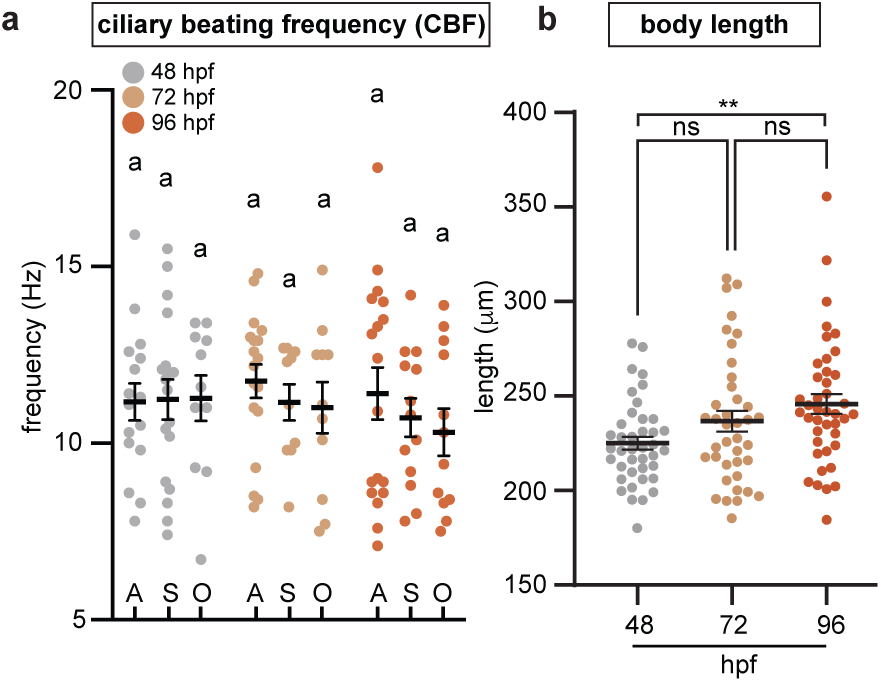
Individual data points for CBF and body length across ages. **a)** Cilia beat at frequencies ranging from 7 to 18 Hz with no significant differences between the means by Kruskal-Wallis test (comparing within region-groups) or mixed-effects analysis (comparing within age-group) with Dunn’s multiple comparisons. **b)** Body length of larvae shows an increasing trend with age; statistical significance by Dunnet’s T3 multiple comparisons. Lines at means ± SEM.

**Figure S5:**
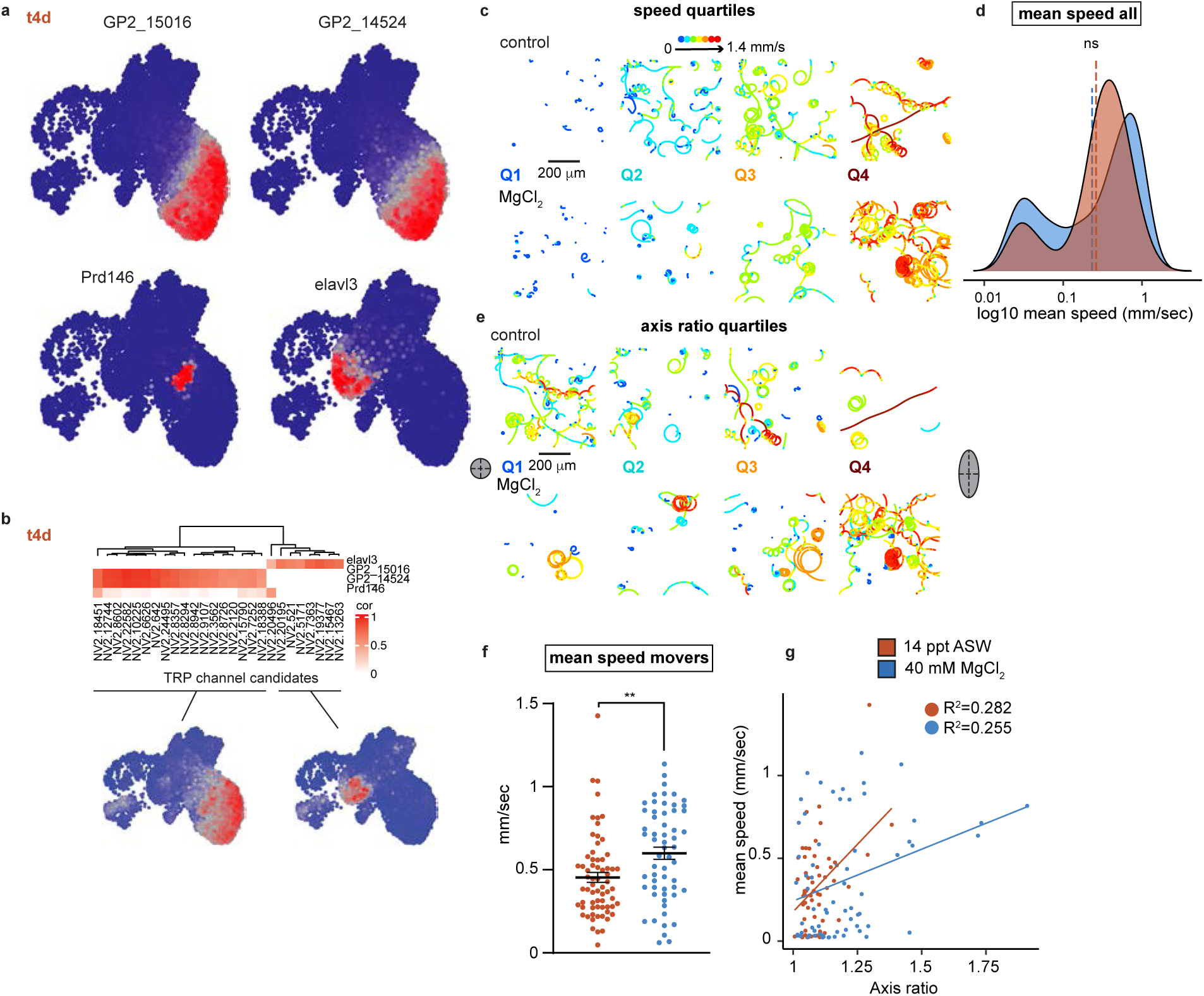
Spatial expression pattern of markers and MgCl_2_ treatment data. **a)** 2D projection of single-cell expression values (right) of marker genes across cell landscape at developmental time t4d. **b)** Correlation between marker expression values (rows) and TRP channel candidates (columns) preferentially expressed in epithelial cells (GP2+) or neuronal cells (elavl3+). 2D cell landscape inserts show projections of average expression values of the corresponding TRP channel candidates. **c)** Tracks of larvae sorted into the defined speed quartiles (Q1-Q4) from the control and MgCl_2_ treated larvae (Ctrl n=84, MgCl_2_ n=88). Colors denote momentary speed. **d)** Distributions of mean speeds between the control and MgCl_2_ treated larvae. Line at the mean, ns p=0.5981 by Mann-Whitney test. **e)** Tracks of larvae sorted into the defined axis ratio quartiles (Q1-Q4) from the control and MgCl_2_treated larvae (Ctrl n=84, MgCl_2_ n=88). Colors denote momentary speed. **f)** Mean speed for moving animals. **p< 0.0015 by Mann-Whitney test (Ctrl n=67, MgCl_2_ n= 58). **g)** Correlation between axis ratio and mean speed for the two treatments.

**Supplementary material: Quantification of analyses, statistical methods and gene IDs related to Figures 1, 2, 3, 4, 5, S3, S4, S5.**

